# Anatomical and functional organization of the human substantia nigra and its connections

**DOI:** 10.1101/120857

**Authors:** Yu Zhang, Kevin Larcher, Bratislav Misic, Alain Dagher

**Affiliations:** Montreal Neurological Institute, McGill University, Montreal, Quebec, Canada.

**Keywords:** parcellation, substantia nigra, dopamine, reward prediction error, value, salience, impulsivity

## Abstract

We investigated the anatomical and functional organization of the human substantia nigra (SN) using diffusion and functional MRI data from the Human Connectome Project. We identified a tripartite connectivity-based parcellation of SN with a limbic, cognitive and motor arrangement. The medial SN connects with limbic striatal and cortical regions and encodes value (greater response to monetary wins than losses during fMRI), while the ventral SN connects with associative regions of cortex and striatum and encodes salience (equal response to wins and losses). The lateral SN connects with somatomotor regions of striatum and cortex and also encodes salience. Connectivity within the value-coding medial SN network was associated with a measure of decisional impulsivity, while the salience-coding ventral SN network was associated with motor impulsivity. In sum, there is anatomical and functional heterogeneity of human SN, and separate SN networks underpin value versus salience coding, and impulsive choice versus impulsive action.

## Introduction

Dopamine innervation to the cerebral hemispheres originates in the substantia nigra (SN) and ventral tegmental area (VTA) of the midbrain. In monkeys, SN/VTA dopamine neurons display variations in both anatomy and function. Anatomically, SN neurons can be divided into three tiers based on their staining, appearance, and connectivity with the striatum (Haber, 2014; Haber and Knutson, 2010): moving from a dorso-medial to ventro-lateral location in midbrain, dopamine neurons project to limbic, associative and then motor striatum. All three subdivisions send dendrites ventrally into the adjacent SN pars reticulata (Haber and Knutson, 2010). Distinct functional characteristics have also been reported for SN/VTA neurons by recording neural activity during appetitive and aversive outcomes (Matsumoto and Hikosaka, 2009; Nomoto et al., 2010). Cells in ventromedial SNc and VTA encode a value signal, being excited by appetitive events and inhibited by aversive events. Neurons in lateral SN may encode a salience signal, responding to both appetitive and aversive stimuli.

Dopamine plays a crucial role in decision-making and reinforcement learning by encoding a reward-prediction error signal (Bayer and Glimcher, 2005; Glimcher, 2011; Schultz et al., 1997). More recently, the role of dopamine in motivational and cognitive processing has been extended by descriptions of responses not only to rewarding outcomes but also to novel, salient, and possibly aversive experiences (Bromberg-Martin et al., 2010; Lisman and Grace, 2005; Matsumoto and Hikosaka, 2009; Redgrave and Gurney, 2006).

An important clinical aspect of dopamine signaling is its role in impulsivity, defined as a tendency to act rapidly and prematurely without appropriate foresight (Dagher and Robbins, 2009; Dalley and Robbins, 2017; Morris and Voon, 2016). Impulsivity is a key feature of drug addiction, obesity, and attention deficit hyperactivity disorder (ADHD). It can be divided into different components (Meda et al., 2009). Decisional impulsivity is characterized by a tendency to make maladaptive or inappropriate choices and is typically tested with the Delay Discounting task. Motor impulsivity, on the other hand, refers to premature responding or an inability to inhibit an inappropriate action, and can be tested using Go/No Go type tasks.

Discrete neural networks may underlie different forms of impulsivity: VS and ventromedial prefrontal cortex (vmPFC), which encode stimulus value, are implicated in decisional impulsivity (Kable and Glimcher, 2007; McClure et al., 2004; Sellitto et al., 2010); somatomotor cortex, supplementary motor area (SMA), inferior frontal gyrus (IFG), anterior insula and dorsal striatum are thought to play a role in motor impulsivity (Bari and Robbins, 2013; Cai et al., 2014; Chikazoe et al., 2009). All of these brain regions are interconnected with SN: they receive dopamine innervation, and send back direct or indirect projections that can modulate dopamine neuron activity (Haber and Knutson, 2010).

In this study, we sought to determine anatomical and functional subdivisions of human SN, their connections with striatal and cortical regions, and their role in value versus salience coding, and in different forms of impulsivity. To date, the in-vivo mapping of SN connectivity in humans has been challenging. The brainstem is prone to artifacts from head movement, pulse and respiration, as well as image distortions during data acquisition. Besides, the SN is a relatively small structure that connects with cortical and striatal regions through dense tracts in the internal capsule (Meola et al., 2016), which causes difficulties for diffusion tractography (Jbabdi et al., 2015). Here we attempt to overcome these limitations using data from the human connectome project (HCP), taking advantage of the high spatial resolution and rich collection of multimodal measures. First, we used diffusion tensor imaging (DTI) to identify subdivisions of SN according to their connectivity patterns with the rest of brain. We then mapped the distinct connectivity profiles for each subdivision. Next, we used an fMRI gambling task to differentiate BOLD activity in SN subdivisions and their projections in terms of responding predominantly to value or salience. Finally, we related individual differences in SN connectivity and activation to performance on two tasks of decisional and motor impulsivity to reveal dissociable neural substrates underlying impulsive choice and impulsive action.

## Materials and Methods

### Subjects and Data Acquisition

Data from 485 healthy individuals (age: 29.1 ± 3.5 years, 202 females) were obtained from the 500-subject release of the Human Connectome Project (HCP) database from March 2015. Participants with body mass index (BMI) lower than 18 were considered as underweight and excluded from this study. Multimodal data used here include structural MRI, diffusion-weighted images, functional MRI during a gambling task and behavioral measures of impulsivity. The scanning procedures are described in detail in (Van Essen et al., 2013) and available online (https://www.humanconnectome.org/documentation/S500/HCP_S500_Release_Reference_Manual.pdf).

### Diffusion MRI

Diffusion data were collected using a single-shot, single refocusing spin-echo, echo-planar imaging sequence with 1.25 mm isotropic spatial resolution (TE/TR = 89.5/5520 ms, FOV = 210 × 180 mm). Three gradient tables of 90 diffusion-weighted directions, and six *b* = 0 images each, were collected with right-to-left and left-to-right phase encoding polarities for each of the three diffusion weightings (*b* = 1000, 2000, and 3000 s/mm^2^). Diffusion data were downloaded in a minimally pre-processed form using the HCP Diffusion pipeline (Glasser et al., 2013) including: normalization of *b*0 image intensity across runs; correction for EPI susceptibility and eddy-current-induced distortions, gradient-nonlinearities and subject motion. Additional pre-processing was performed using BEDPOSTX from the FSL Diffusion Toolbox (Behrens et al., 2007) to estimate probability distributions for at most three fiber directions at each voxel. A T1-weighted image (FOV = 224 mm, matrix = 320, 256 slices; TR = 2400 ms, TE = 2.14 ms, TI = 1000 ms, FA = 8°; 0.7 mm isotropic resolution) registered into the diffusion space was also provided for each subject, which was employed for the nonlinear registration of the SN seed from MNI standard space (http://www.bic.mni.mcgill.ca/ServicesAtlases/ICBM152NLin2009) to individual diffusion space using FNIRT from the FSL package. In total, 430 subjects’ data were pre-processed and passed quality control. The parcellation of SN was carried out on 120 randomly selected subjects in order to limit computation time and data storage. To test the robustness of parcellation, we randomly divided the 120-subject dataset into two and applied the parcellation procedure (as explained below) independently in each group. Specifically, the first 60 subjects were used as the test group to reveal the underlying organizational pattern of SN. A second group of 60 subjects was then used as the replication group to test the stability of our parcellation maps. All diffusion data (N=430) were then used during the tractography analysis to map the connectivity profiles of each of the SN subdivisions identified by parcellation.

### Functional MRI with gambling task

HCP fMRI data were acquired at a high-resolution (2 mm isotropic) with a 32 channel head coil on a modified 3T Siemens Skyra using fast temporal sampling and multiband pulse sequences (TR=720 ms, time to echo = 33.1 ms, flip angle = 52°, bandwidth = 2,290 Hz/pixel, in-plane field of view = 208 × 180 mm, 72 slices, and 2.0 mm isotropic voxels, with a multiband acceleration factor of 8) (Uğurbil et al., 2013; Van Essen et al., 2013). During the gambling task (Barch et al., 2013), participants were asked to guess the number on a mystery card that displayed a question mark. The card numbers ranged from 1 to 9 and participants were asked to guess whether the mystery card number was above or below 5 by pressing one of two buttons. Feedback was the actual number on the card along with the outcome of the trial: 1) a green up arrow with “$1” for reward trials, 2) a red down arrow with “-$0.50” for loss trials; or 3) the number 5 and a gray double headed arrow for neutral trials. The cue with the mystery card was presented for up to 1.5 s, followed by 1.0 s of feedback. Before starting the next trial, there was a 1.0 s ITI with a “+” presented on the screen. All participants received their winnings after completing the task. The fMRI data were downloaded in a pre-processed form performed using the HCP fMRIVolume pipeline (Glasser et al., which included motion correction, field-map correction, nonlinear registration into MNI standard space and spatial smoothing using a Gaussian kernel of FWHM = 4 mm in the volume space.

### Impulsivity measures

Measures from two behavioral tests of impulsivity were used for each subject: the Delay Discounting task and the Flanker Inhibitory Control Task. Delay discounting was selected as the measure of impulsive choice. It describes the temporal discounting of monetary rewards, in which subjects choose between a smaller immediate and a larger delayed reward. An adjusting-amount approach was used for the delay discounting paradigm (Estle et al., 2006), in which delays are fixed and reward amounts are adjusted on a trial-by-trial basis based on participants' choices. The indifference point for each delay period is first identified as the monetary amount (for the delayed reward) at which the participant is equally likely to choose between the immediate and delayed reward. The indifference points indicate the subjective value of delayed outcomes. By plotting indifference points for each delay, a discounting measure of area-under-the-curve (AUC; (Myerson et al., 2001)) was calculated by first normalizing the indifference points by the maximum value for all delay periods and then summing the areas underneath the curve using the following equation: (x_2_ – x_1_) *(y_1_ + y_2_)/2, where x_1_ and x_2_ are successive delays and y_1_ and y_2_ are the indifference points associated with those delays. The AUC value ranges from 1 (no discounting) to 0 (maximum discounting) with larger values representing less impulsive decisions. Only the high monetary amount ($40,000) was used here considering that its AUC is approximately uniformly distributed across all subjects. The Flanker Inhibitory Control Task from the NIH Toolbox (http://www.nihtoolbox.org) was selected as a measure of response inhibition. During the Flanker task (Eriksen and Eriksen, 1974), participants are required to indicate the left–right orientation of a centrally presented arrow while inhibiting their attention to the surrounding flanking stimuli. There are two types of trials, one with the middle arrow pointing in the same direction as the "flankers" (congruent) and the other with it pointing in the opposite direction (incongruent). A two-vector scoring method (Zelazo et al., 2014) was employed to combine the accuracy (range from 0 to 5 with the lower values representing fewer correct responses on both congruent and incongruent trials), and rescaled reaction time (range from 0 to 5 with the lower values representing slower reaction time). For individuals with low accuracy levels (less than 80%), only the accuracy score was calculated, while both reaction time score and accuracy score were combined when the accuracy level was greater than 80%. The final scores were additionally adjusted for age. Higher scores represent both higher accuracy levels and faster reaction times, and therefore better inhibitory control.

### Seed regions

A mask of substantia nigra was generated from a 7T MRI atlas of basal ganglia based on high-resolution MP2RAGE and FLASH scans (Keuken and Forstmann, 2015). The authors manually delineated the main structures of basal ganglia in 30 young healthy participants (age: 24.2 ± 2.4 years, 14 females) to generate a probabilistic atlas for each structure. The entire region of SN (Figure 2A) was extracted from the probabilistic atlas with a threshold of 33% of the population (i.e. retaining voxels labeled as SN in at least 10 out of 30 subjects) yielding masks of volume equal to approximately 300 mm^3^ in each hemisphere.

Other regions of interest of brain areas involved in reward and salience processing were defined as follows. Ventral striatum and ventral medial prefrontal cortex (vmPFC) were defined by drawing a 6-mm sphere around the peak coordinates from a fMRI meta-analysis of subjective value (Bartra et al., 2013). Two salience-related areas, the dorsal anterior cingulate cortex (dACC) and anterior insular cortex, were defined by drawing a 6-mm sphere around the peak coordinates of the salience network identified from resting state fMRI (Seeley et al., 2007). The MNI coordinates of these regions of interest are listed in Table 1. Here, we hypothesized that ventral striatum and vmPFC were part of a value-coding system (with greater activation to reward than punishment), while dACC and anterior insula were more involved in salience-coding (i.e. responding similarly to reward and punishment).

**Table 1.** Regions of interest used in this study and their BOLD activity during the gambling task.

| Brain Regions | *x* | *y* | *z* | *Brain Activity (T-score)* |  |  |
| --- | --- | --- | --- | --- | --- | --- |
|  |  |  |  | *Reward* | *Punishment* | *RPE* |
| SN subdivisions |  |  |  |  |  |  |
| vSN | ± 9 | -13 | -12 | 11.12 ** | 8.63 ** | 1.85 |
| medial SNc | ± 8 | -16 | -12 | 17.45 ** | 12.48 ** | 3.96 ** |
| lateral SNc | ± 12 | -17 | -9 | 9.72 ** | 8.87 ** | 0.93 |
| Ventral Striatum | ± 12 | 15 | -6 | 8.91 ** | - 9.33 ** | 16.25 ** |
| vmPFC | ± 6 | 45 | -9 | - 26.72 ** | - 32.37 ** | 8.75 ** |
| Anterior insula | ± 32 | 22 | -6 | 35.54 ** | 36.62 ** | 1.07 |
| dACC | ± 4 | 40 | 24 | 8.43 ** | 7.64 ** | 1.37 |

Notes: \*\* : p-value< 0.01; \* : p-value< 0.05 with FDR correction
SN: substantia nigra; vSN: ventral subregion of SN; SNc: SN pars compacta; vmPFC: ventral medial prefrontal cortex; dACC: dorsal anterior cingulate cortex.

### Connectivity-based parcellation of SN

A data-driven connectivity-based brain parcellation procedure was used (Figure 1, also described in (Fan et al., 2016). First, probabilistic tractography was applied by sampling 5000 streamlines at each voxel within the seed mask of SN. The whole-brain connectivity profile for each voxel was saved as a connectivity map, where the intensity shows how many streamlines reach the target area and is therefore a measure of the connectivity strength between the seed and target. These connectivity maps were used to generate a connectivity matrix with each row representing the whole-brain connectivity profile of one seed voxel. Next, a correlation matrix was calculated as a measure of similarity between the connectivity profiles of each voxel pair (Johansen-Berg et al., 2004). Then, spectral clustering (Shi and Malik, 2000) was applied to the similarity matrix to identify clusters with distinct connectivity profiles. We applied this procedure separately for each subject and each hemisphere to generate a series of parcellation maps for all individuals at different resolutions (i.e. numbers of regions/parcels). Here, considering the small size of the SN, we chose cluster numbers ranging from 2 to 8 in each hemisphere and chose the most stable and consistent parcellation map (see below).

**Figure 1.**
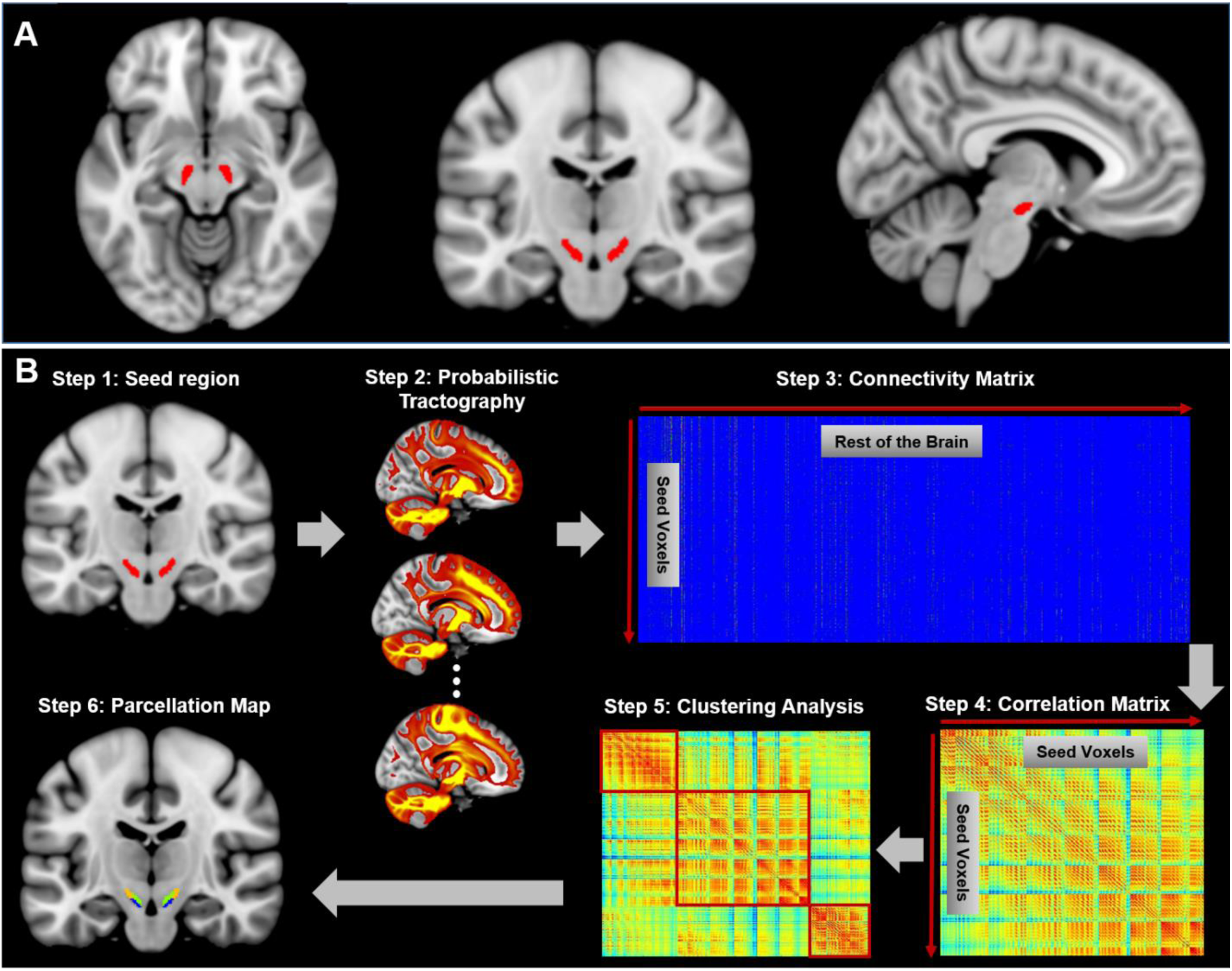
Connectivity-based brain parcellation procedure. After defining the seed region (step 1), probabilistic tractography was applied by sampling 5000 streamlines at each voxel within the seed mask (step 2). Then, these whole-brain connectivity profiles were loaded into a connectivity matrix with each row representing the connectivity profile of each seed voxel (step 3). Next, a correlation matrix was calculated as a measure of similarity between seed voxels (step 4). Then, spectral clustering was applied to the similarity matrix (step 5) and multiple subdivisions were identified within the seed region (step 6). The entire procedure was applied independently for each hemisphere and each subject.

An additional group-parcellation procedure was applied to summarize the general pattern of parcellation across subjects. Specifically, a consensus matrix *S* was defined based on each individual parcellation map, with each element *S_ij_* = 1 if and only if voxel *v_i_* and voxel *v_j_* belong to the same cluster. Then, a group consensus matrix was generated by averaging the consensus matrices from all subjects. The final group parcellation map was generated by performing spectral clustering again on the group consensus matrix (Fan et al., 2016; Zhang et al., 2015).

The optimum parcellation solution (i.e. number of parcels) was determined by evaluating the reproducibility of parcellation maps through a split-half procedure. Specifically, we randomly split the entire group into two non-overlapping subgroups 100 times and generated the group parcellation maps for each subgroup separately. The consistency between each pair of parcellation maps was evaluated by different stability indices, including normalized mutual information (NMI) (Zhang et al., 2015), Dice coefficient (Zhang et al., 2014) and Cramer’s V (Fan et al., 2014). The average indices among 100 samples were calculated to represent the stability of each parcellation. The suitable cluster number was then determined by searching for the local peaks in the stability curve. In addition, topological similarity of the parcellation solutions between the two hemispheres was also calculated as a measure of stability (Fan et al., 2016).

### Connectivity profile of each subdivision

Based on the obtained parcellation map of SN, we mapped the anatomical connectivity profiles of each subdivision by performing probabilistic tractography with 10,000 streamlines from each subarea. The resulting connectivity maps were first normalized by the size of seed region and total number of streamlines (i.e. 10,000) in order to generate the relative tracing strength from the seed to the rest of the brain. A threshold of 0.001 (i.e. 10 out of 10,000) was then used to remove noise effects of fiber tracking. The resulting individual tractograms were combined to generate a population map of the major fiber projections for each SN subdivision. Another probabilistic threshold of 50% was applied to the population fiber-tract maps (i.e. at least half of subjects showing each retained fiber tract). This resulted in a group averaged tractogram for each subdivision of SN. Finally, a maximum probability map (MPM) of fiber tracts, which represents distinct components of SN projections derived from each subdivision, was also generated based on the population fiber-tract maps. Specifically, a connectome mask was first generated for each subdivision by binarizing its group tractography map with connectivity probability at 0.01. Note that different probability thresholds do not change the organizational pattern, but only enlarge or shrink the coverage of major fiber tracts (Figure 4-Figure Supplement 1). Each voxel within the combined connectome mask was then classified according to the SN subdivisions with which it had the highest connectivity. This calculation of MPM on probabilistic tractography has been widely used in subdividing brain structures, including thalamus (Behrens et al., 2003), amygdala (Saygin et al., 2011) and striatum (Cohen et al., 2009). Here, we use this method to generate the organizational topography of fiber projections among SN subdivisions.

A quantitative representation of the connectivity profiles was also generated by calculating the connectivity fingerprints between each SN subdivision and each cortical/subcortical area. A recently published brain atlas based on anatomical connectivity profiles (Fan et al., 2016) was chosen to define the target areas, consisting of a fine-grained parcellation of frontal, parietal, temporal, occipital cortex, limbic areas, as well as striatum and thalamus. The relative connectivity strength between each SN subdivision and each brain parcel was calculated by averaging the tractography values of the population fiber-tract maps. An additional normalization step was applied by dividing the tractography values with the total number of streamlines among the three SN subdivisions. These normalized connectivity values were then used to estimate the connectivity fingerprints for each SN subdivision.

### Neural activity during Gambling task and correlations with impulsivity measures

The event-related gambling task fMRI paradigm from HCP was chosen to identify brain areas responding to monetary reward and salience. Specifically, a general linear model (GLM) implemented in FSL's FILM (Woolrich et al., 2001) was used to estimate the neural activity during the feedback of reward (i.e. winning $1), punishment (i.e. losing $0.5) or neutral outcomes (i.e. no gain or loss) by convolving each event epoch with a double gamma “canonical” hemodynamic response function. The temporal derivatives of the events were also added in the model as confounds. Next, three types of contrasts were defined: wins minus neutral, losses minus neutral, and wins minus losses. It is worth mentioning that, in this task design, the difference in BOLD response to reward and punishment is proportional to the reward prediction error (RPE) -related signal (i.e. the expected value on each trial equals $1 * 0.5 + (−$0.5) * 0.5 = $0.25; prediction error for win trials equals $1 − $0.25 = $0.75; prediction error for loss trials equals (−$0.5) − $0.25 = −$0.75; thus the prediction error is $0.75 for win trials and −$0.75 for loss trials). The resulting parameter estimates (cope) of contrast images from the two acquisitions using different phase encoding directions (i.e. LR and RL coding) were combined to generate individual BOLD activity during win and loss outcomes (Barch et al., 2013). The final effect size of each contrast was extracted for each subject and each region of interest, including SN subdivisions, VS, vmPFC, dACC and anterior insula, and was remodeled into effects of value-coding (i.e. difference in response to reward and punishment) and salience-coding (i.e. averaged response to reward and punishment). Moreover, to examine the association between task-related BOLD activity during rewarding and aversive outcomes and behavioral impulsivity measures, a correlation analysis was performed between neural activity under value- or salience-coding conditions and decisional impulsivity (measured by delay discounting task) or motor impulsivity (measured by the Flanker inhibitory control task), with age and gender as covariates of no interest.

## Results

### Subdivision of Substantia Nigra

Three stable subdivisions were identified in the substantia nigra of each hemisphere (Figure 2B): a dorsolateral area corresponding to lateral part of SN pars compacta (lateral SNc – hereafter: lSNc), a dorsomedial area corresponding to medial part of SN pars compacta (medial SNc - mSNc) and a ventral area (vSN). The MNI coordinates for the center of mass of each SN subdivision are listed in Table 1. The three subdivisions were of similar volume on average (vSN: 93/90 *mm*^3^ for left/right; mSNc: 90/117 *mm*^3^ for left/right; lSNc: 117/98 *mm*^3^ for left/right). This separation of dorsomedial, lateral and ventral SN coincides with descriptions of dorsal, middle and ventral tiers of midbrain dopamine cells in primates (Figure 2-Figure Supplement 1), which have distinct afferent and efferent striatal and cortical projections (Haber, 2014; Haber and Knutson, 2010).

**Figure 2.**
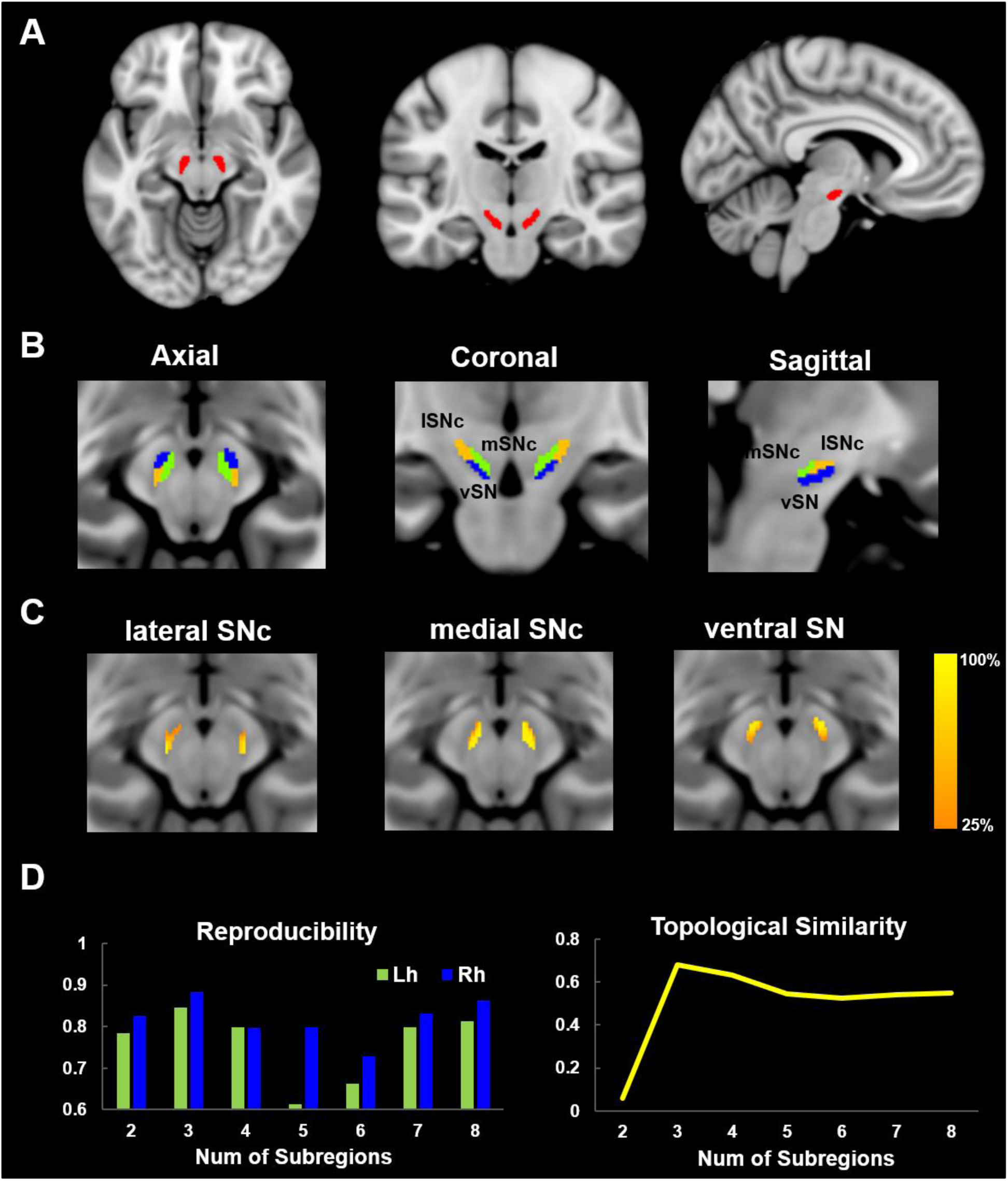
Parcellation of Substantia Nigra based on anatomical connectivity profiles. A) Definition of the seed region. Substantia Nigra was extracted from a 7T atlas of Basal ganglia based on high-resolution MP2RAGE and FLASH scans (Keuken and Forstmann, 2015). B) Parcellation map of SN on 60 healthy young subjects. SN was subdivided into three subregions: a dorsolateral area corresponding to lateral part of SN pars compacta (lSNc), a dorsomedial area corresponding to medial part of SNc (mSNc) and a ventral area (vSN). C) Probabilistic map of each SN subdivision. D) Reproducibility and topological similarity of SN parcellation. The three-cluster parcellation of SN showed both high reproducibility, as assessed by repeated split-half resampling (mean NMI = 0.85 and 0.88, respectively for left and right SN) and high inter-hemispheric topological similarity (mean NMI=0.68).

The optimum number of subregions within SN was determined by evaluating both reproducibility of parcellation using repeated split-half resampling and topological similarity across hemispheres. As shown in Figure 2D, the three-subdivision parcellation of SN showed both high reproducibility (mean NMI = 0.85 and 0.88, respectively for left and right SN) and high inter-hemispheric topological similarity (mean NMI=0.68). The same conclusion was drawn from the other stability indices, the Dice coefficient and Cramer’s V (Figure 2-Figure Supplement 3). Moreover, the stability of the parcellation is also supported by the probabilistic maps of each subdivision (Figure 2C), where the intensity represents the probability of subdivision assignment over the population at each SN voxel. Finally, the replication of the parcellation procedure on a second group of 60 randomly selected subjects from the HCP dataset yielded very similar results (NMI=0.95; Figure 2-Figure Supplement 2).

### Connectivity patterns of SN subdivisions

Distinct connectivity profiles were identified for each subdivision of SN by performing probabilistic fiber tractography from each subdivision (Figure 3) on 430 HCP datasets. Specifically, the dorsolateral subregion (i.e. lateral SNc) mainly connected with the somatic motor and sensory cortex in pre-/post-central gryus; the dorsomedial subregion (i.e. medial SNc) showed dominant connections to limbic regions including lateral and medial OFC, hippocampus and amygdala. The ventral subregion (i.e. vSN) preferentially connected to prefrontal cortex, anterior cingulate cortex and anterior insula. These connectivity maps reveal a limbic-cognitive-motor organizational topography of SN fiber projections.

**Figure 3.**
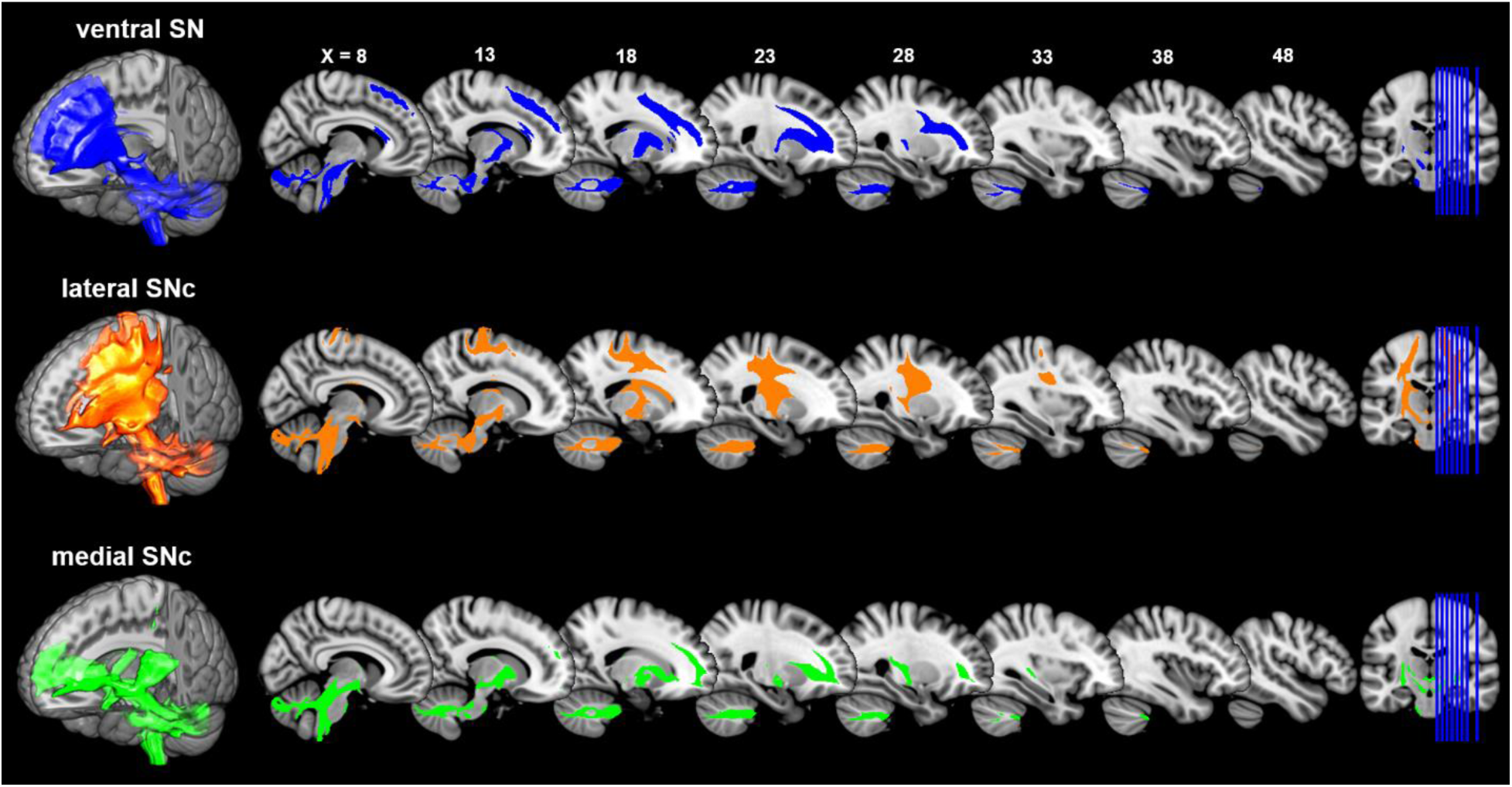
Connectivity patterns of the subdivisions of Substantia Nigra. Probabilistic fiber tractrography was performed for each SN subdivision to map their whole-brain connectivity patterns. The population tract maps are shown with a threshold of connectivity probability at 0.1 and rendered by using MRIcron on the ICBM152 brain template.

This limbic-cognitive-motor topology of SN projections was also evident from the MPM of the tractograms (Figure 4 and Figure 4-Figure Supplement 1). Particularly in prefrontal cortex, a clear rostocaudual pattern of SN projections was present, with medial SNc mainly projecting to the most rostral part including OFC and frontal pole; vSN connecting to lateral and dorsomedial prefrontal cortex including middle frontal gyrus and inferior frontal gyrus; and lateral SNc showing connections with the sensorimotor and somatosensory cortex. A similar rostrocaudal distribution of fiber tracts was also seen in striatum, with medial SNc mainly connecting with ventral striatum, vSN connecting via the anterior limb of the internal capsule with the body of caudate and anterior part of putamen (associative striatum), and lateral SNc connecting via the posterior limb of the internal capsule with the posterior part of putamen (motor striatum). Moreover, vSN strongly connected with the external part of the globus pallidus while medial SNc showed dominant connections with the internal part of the globus pallidus and ventral pallidum.

**Figure 4.**
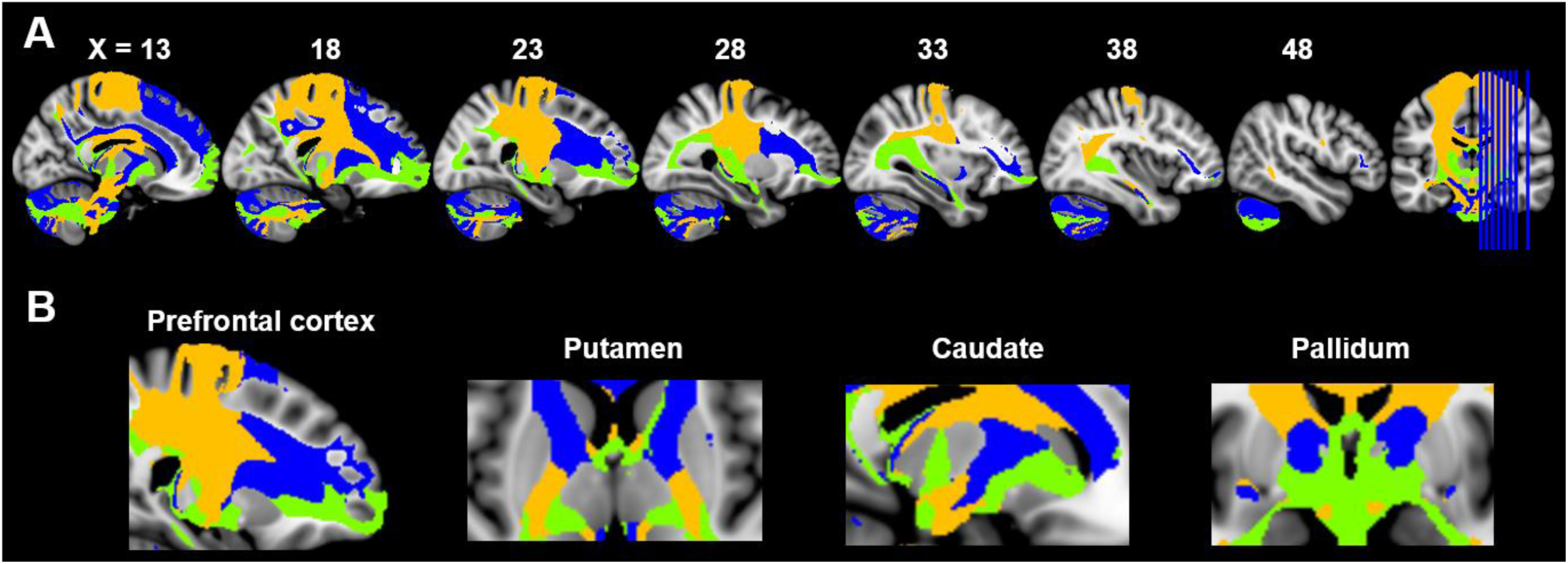
Maximum probability tractograms of the subdivisions of Substantia Nigra. A limbic-cognitive-motor organizational topography of SN projections is shown in multi-slice views (A) and particularly for prefrontal cortex, striatum and pallidum (B). MPM tractograms were generated by assigning each voxel to the corresponding SN subdivision with which it showed the greatest connections. SN subdivisions: vSN (blue), mSNc (green) and lSNc (orange).

A similar organizational pattern was also revealed by analyzing the SN connections to seven canonical resting-sate networks (Yeo et al., 2011), with clear dissociations of fiber projections among the three SN subdivisions (Figure 5-Figure Supplement 2). Specifically, medial SNc preferentially connected to the limbic and visual networks; vSN dominantly connected to the frontoparietal and default-mode network; and lateral SNc mainly connected to the somatomotor and dorsal attention networks.

**Figure 5.**
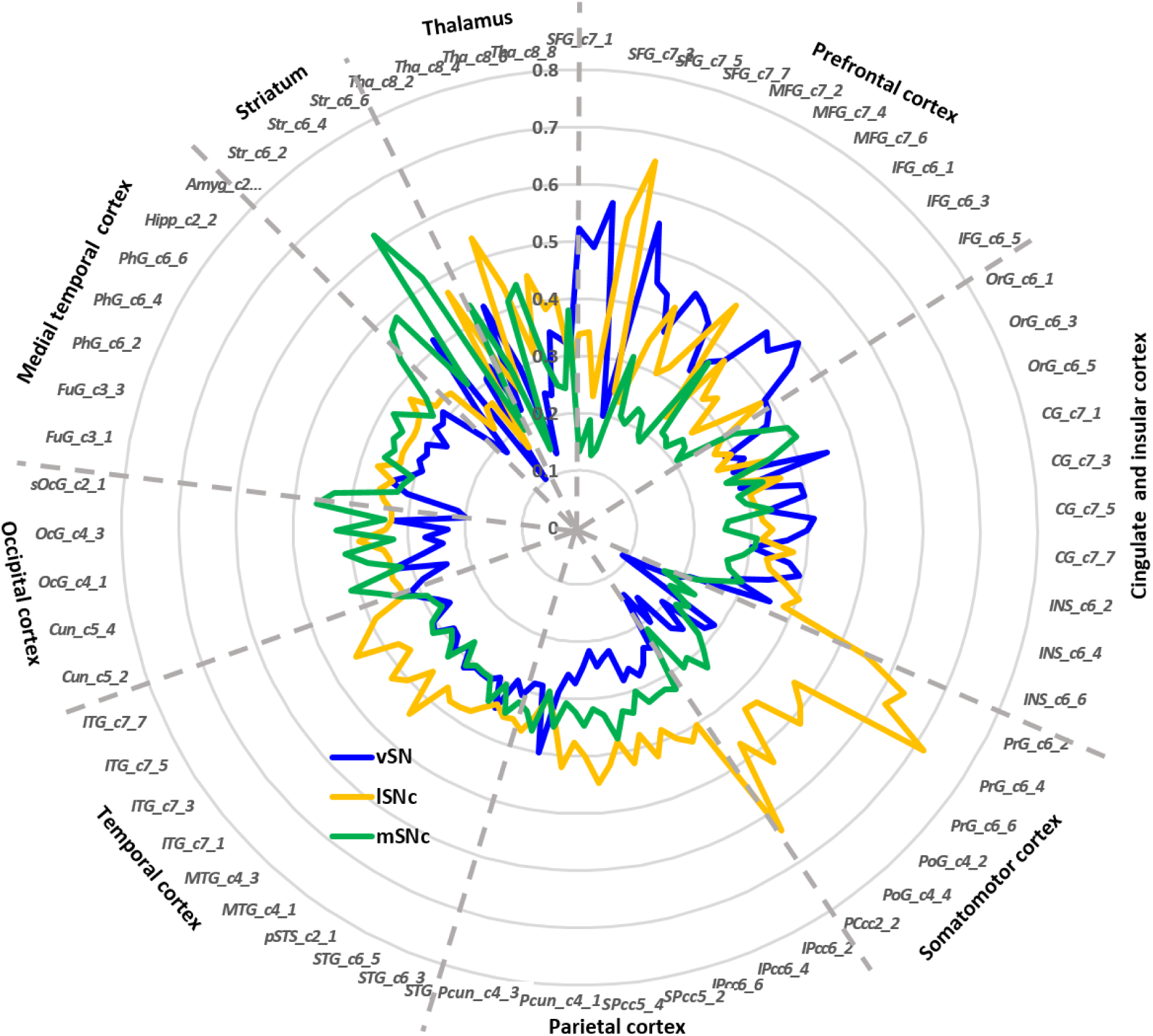
Connectivity fingerprints of the subdivisions of Substantia Nigra. The connectivity fingerprints of SN subdivisions were calculated based on a whole-brain atlas (Fan et al., 2016). The relative connectivity strength between each target (i.e. parcels in the brain atlas) and each SN subdivision is plotted. An organizational topography of SN projections emerges with vSN strongly connected to prefrontal cortex, lateral SNc mostly connected with sensorimotor cortex, and medial SNc to limbic and striatal regions.

Finally, connectivity fingerprints of the three SN subdivisions were generated by mapping the connectivity profiles to a fine-grained whole brain anatomical connectivity atlas (Fan et al., 2016). As shown in Figure 5, the three subdivisions of SN showed distinct connectivity profiles in frontal, parietal, temporal and subcortical areas. Specifically, most prefrontal areas showed the strongest connections to vSN (Figure 5-Figure Supplement 1A), except for several subregions in superior and middle frontal gyrus, for instance lateral and medial area 6 (i.e. areas SFG_c7_4/5 and MFG_c7_6), more strongly connecting to lateral SNc. The somatic motor and sensory cortex preferentially connected with lateral SNc (Figure 5-Figure Supplement 1B). Meanwhile, most limbic and striatal regions were targeted by fiber tracts derived from medial SNc (Figure 5-Figure Supplement 1C), except for the tail of caudate (i.e. area Str_c6_5) showing relatively stronger connections with lateral SNc and anterior putamen (i.e. area Str_c6_2) showing higher connections to vSN.

In summary, all of the connectivity profiles of the SN subdivisions are consistent with a limbic (medial SNc), motor (lateral SNc), and cognitive (vSN) functional organization.

### Brain activity during gambling task

In order to ascertain different functional roles of SN subdivisions, we explored their BOLD response to rewarding and aversive stimuli during the fMRI gambling task. Brain activation maps of value-coding (i.e. contrast of the difference in response to reward versus punishment) and salience-coding (i.e. contrast of the mean response to reward and punishment) were assessed by whole-brain analysis using one-sample t-tests and corrected for multiple comparisons using the threshold-free cluster enhancement method (Smith and Nichols, 2009). Significant value-coding was detected in ventral striatum and vmPFC, while salience signals were found in anterior insula, dorsal ACC and dorsal striatum (Figure 6-Figure Supplement 1). Next, specific analysis was performed on the seven regions of interest consisting of the three SN subdivisions, ventral striatum, vmPFC, anterior insula and dorsal ACC. As shown in Figure 6, all three SN subregions were activated by both reward and punishment, but only medial SNc showed significant difference in BOLD response to the two, i.e. significantly greater neural activity to monetary gains than losses (T= 3.96, p< 0.0001). Among the other regions of interest, ventral striatum showed a unique bi-directional pattern, e. strongly activated during reward (T= 8.91, p< 0.0001) and deactivated during punishment (T= −9.33, p< 0.0001). Meanwhile, as a core region in the default mode network (DMN; (Buckner et al., 2008; Raichle, 2015)), vmPFC was deactivated in both conditions (T= −26.72 and −32.37 with p< 0.0001, respectively for reward and punishment). Both ventral striatum and vmPFC showed significantly greater response to rewarding than aversive outcomes (T= 16.25, p< 0.0001 for ventral striatum, T= 8.75, p< 0.00001 for vmPFC). On the contrary, as the core areas of the salience network (Seeley et al., 2007), dACC and anterior insula were activated by both types of trials, with no difference in response to reward and punishment (T=1.37, p= 0.17 for dACC, T= 1.07, p= 0.28 for anterior insula). These results suggest that there are at least two separate brain dopaminergic systems involved during gambling outcomes, with one encoding value signals (i.e. different response to reward and punishment) and the other encoding motivational salience signals (i.e. reacting similarly to rewarding and aversive outcomes).

**Figure 6.**
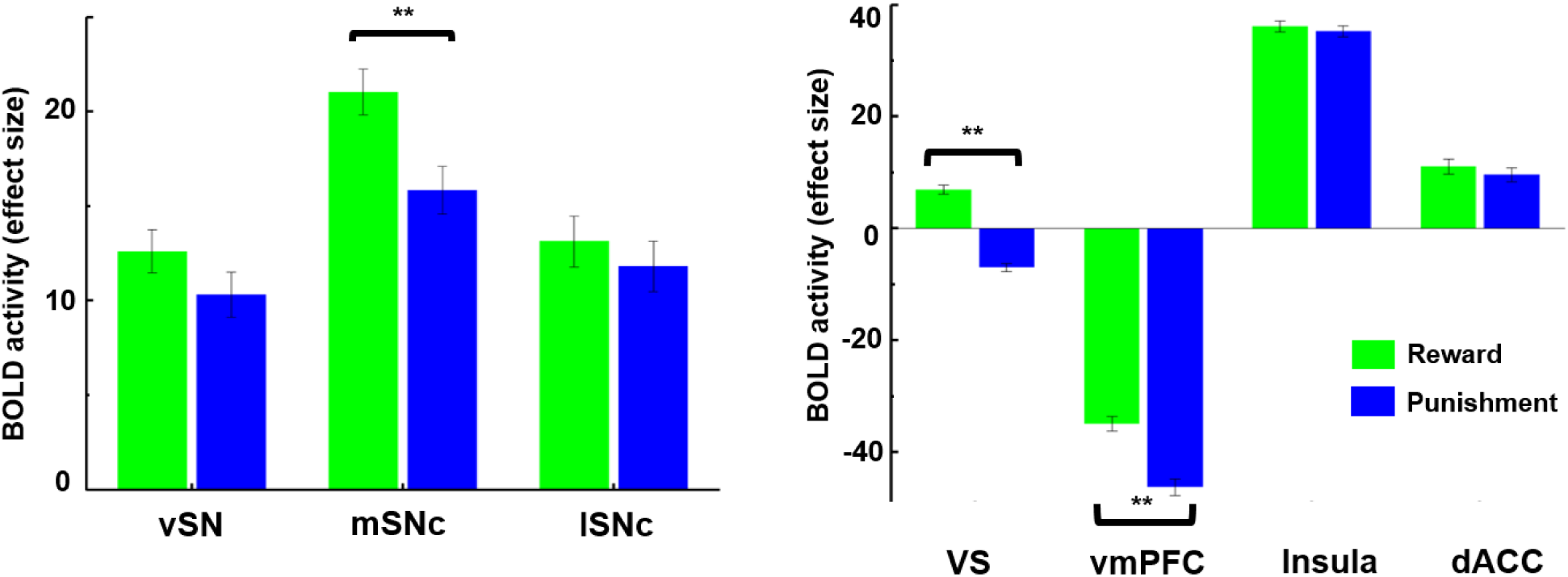
Brain activity in response to rewarding and aversive outcomes in the fMRI gambling task. Among SN subdivisions, only medial SNc showed a significant difference in response to reward and punishment (p< 0.001). The ventral striatum (VS) and ventromedial prefrontal cortex (vmPFC) also responded differently to reward and punishment, with greater BOLD activity to rewarding than aversive stimuli (p< 0.001). Meanwhile, anterior insula and dorsal anterior cingulate cortex (dACC) showed no difference in response to reward and punishment.

### Correlation analysis between brain activity and impulsivity measures

We next sought to determine if brain activity in the value and salience coding system was related to two impulsive traits: decisional and motor impulsivity. Based on the above brain connectivity (Figures 3-5) and activity analyses (Figure 6), we take the value-coding system to consist of mesolimbic pathways projecting between medial SNc and ventral striatum and vmPFC, and the salience-coding system of mesocortical pathways connecting vSN with dACC and anterior insula. We correlated BOLD activity and anatomical connectivity of these brain areas with behavioral measures of decisional and motor impulsivity measures. We reasoned that decisional impulsivity would implicate the value-coding dopamine system, while motor impulsivity would implicate salience or motor system projections.

First, to determine the association between BOLD activity and impulsive behaviors, we performed the correlation analysis between value- and salience-related BOLD response and measures of the two aspects of impulsivity, i.e. the Delay Discounting task for decisional impulsivity and the Flanker inhibitory control task for motor impulsivity. Regarding decisional impulsivity (Figure 7A), the value-related neural activity (i.e. difference in response to monetary gains and losses) in medial SNc and ventral striatum showed a significant negative correlation with the AUC of Delay Discounting (r= −0.1112, p= 0.0164 for medial SNc; r= −0.1029, p= 0.0251 for ventral striatum) but not with Flanker inhibitory control scores (r= −0.0661, p= 0.15 for mSNc; r= 0.0089, p= 0.84 for VS). This indicates that participants with stronger value-coding signals in the putative value-coding areas tend to make more impulsive choices during delay-discounting decisions (i.e. stronger preference for immediate monetary rewards). On the other hand, regarding motor impulsivity (Figure 7B), brain activity related to salience signals (i.e. average response to monetary gains and losses) in dACC and anterior insula showed a significant positive correlation with inhibitory control scores from the Flanker task (r= 0.1078, p= 0.0195 for dACC; r= 0.1296, p= 0.0047 for anterior insula) but not with AUC of Delay Discounting (r= 0.0411, p= 0.37 for dACC; r= −0.0200, p= 0.66 for anterior insula). This indicates that subjects with stronger salience-coding activity in the salience network showed greater capacity for inhibitory control. The relationships were somewhat different for dACC and anterior insula (Figure 7-Figure Supplement 1). Specifically, better inhibitory control (or less motor impulsivity) was associated with greater activity in anterior insula in response to both rewarding and aversive stimuli (r= 0.1169 and 0.1256, p= 0.0108 and 0.0063, respectively for reward and punishment), but only to rewarding outcomes in dACC (r= 0.1550, p= 0.0008).

**Figure 7.**
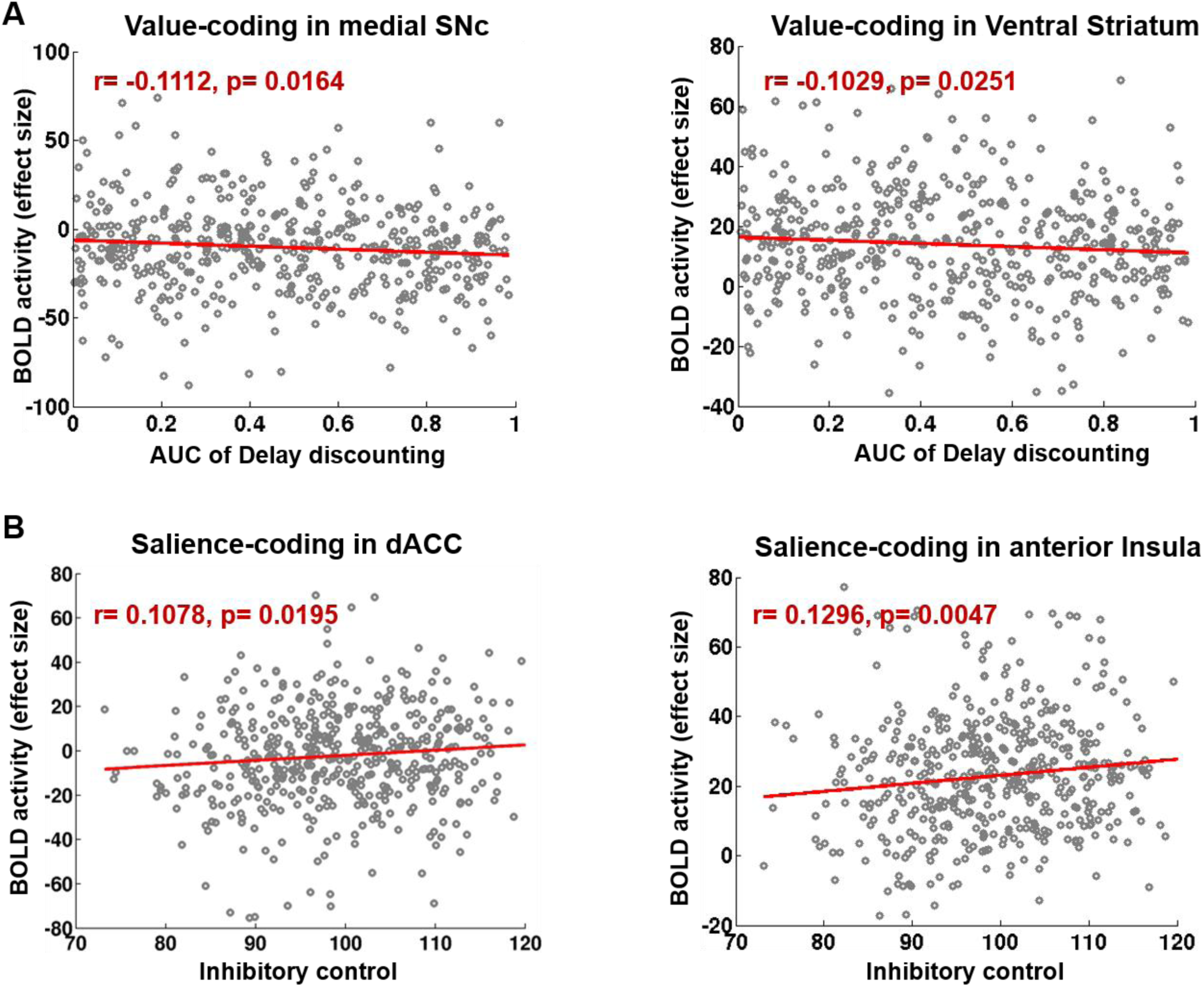
Correlation between value- and salience-coding BOLD activity and behavioral impulsivity measures. Value-related BOLD activity was measured by differences in brain response to rewarding and aversive outcomes, and salience-related BOLD activity was measured by averaged brain response to rewards and penalties. Value-coding activity in mSNc and VS was correlated with the AUC measure of delay discounting task (A), while salience-coding activity in dACC and anterior insula were correlated with the inhibitory control scores of Flanker task (B). Note: greater AUC indicates less impulsivity. (AUC: area under the curve).

### Correlation analysis between anatomical connectivity and impulsivity measures

Associations between the underlying anatomical connections of dopamine pathways and behavioral impulsivity were explored by using Spearman’s Rank-Order correlation analysis. Consistent with the previously described connectivity analysis (Figures 3-5), the mesolimbic pathways terminating in ventral striatum predominantly originated from mSNc, while the mesocortical pathways targeting the salience areas dACC and anterior insula were preferentially derived from vSN (Figure 8A). Moreover, the anatomical connectivity strength between vSN and dACC measured by probabilistic tractography was correlated with the inhibitory control scores from the Flanker task (r= 0.1034, p= 0.03) but not AUC of Delay Discounting (r= 0.0397, p= 0.41), meaning that greater connectivity in the mesocortical pathway was associated with better inhibitory control (less impulsivity). Meanwhile, the anatomical connectivity strength from mSNc to vmPFC showed a significant correlation with the AUC measure of Delay Discounting task (r= 0.122, p= 0.01) but not Flanker inhibitory control scores (r= −0.0224, p= 0.64), meaning that greater connectivity in the value-coding system was associated with lower decisional impulsivity (greater AUC means lower temporal discounting). It is worth mentioning that SN projections targeting vmPFC were equally contributed to by all three subdivisions, which may suggest that, as a connectional hub, vmPFC integrates distributed information to support the valuation process during decision-making (Benoit et al., 2014; Roy et al., 2012).

**Figure 8.**
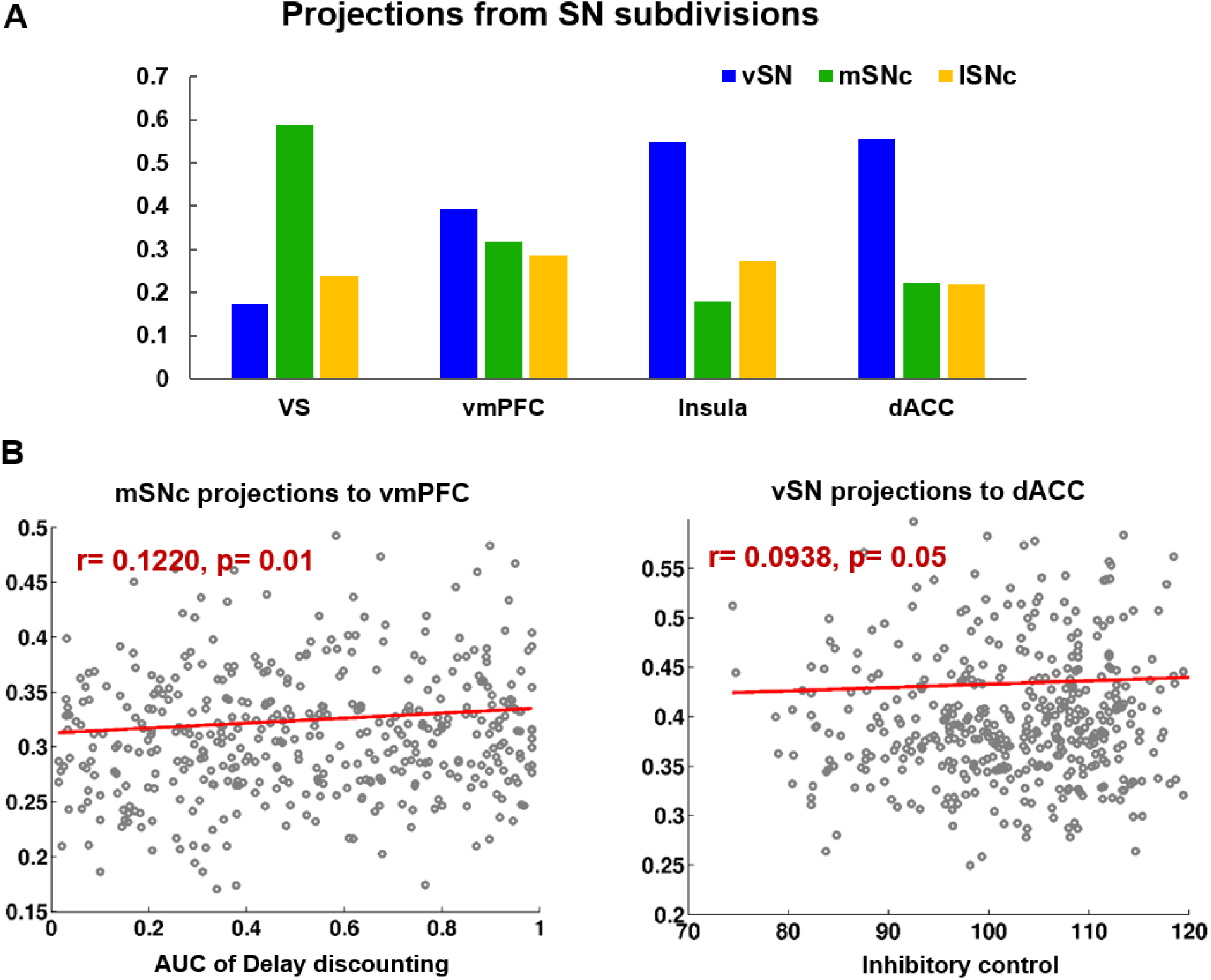
Correlation between DTI projections from SN and impulsivity measures. Two types of SN projections were revealed by the probabilistic tractography results (A), with VS mainly connecting to mSNc, while dACC and anterior insula connecting with vSN. B) Correlation analysis between the two SN projections and the decisional impulsivity (measured by the delay discounting task) and motor impulsivity (measured by the Flanker inhibitory control task).

## Discussion

### Subdivisions of SN

We used a connectivity-based parcellation scheme to subdivide human SN based on its anatomical connectivity profile with the rest of the brain. A tripartite pattern of SN was revealed, consisting of a medial (mSNc), a lateral (lSNc) and a ventral (vSN) tier. A similar anatomical and connectional differentiation of SN has been widely described in monkeys. Indeed, many studies report that midbrain dopamine neurons can be divided into two or three tiers (François et al., 1999; Haber and Knutson, 2010; Lynd-Balta and Haber, 1994), with a dorsal calbindin-positive tier that extends medially to the VTA, and a ventral calbindin-negative tier whose dendrites extend ventrally into the pars reticulata of the substantia nigra. This ventral tier can be further subdivided into a more medio-dorsal densocellular group and a ventro-lateral group of columnar cells (Haber, 2014). Tracer studies in monkeys have been used to map the striatal afferent and efferent projections of these SN subdivisions (Haber et al., 2000; Haber and Knutson, 2010; Lynd-Balta and Haber, 1994). The dorsal tier mainly connects with ventromedial striatum, while the ventral tiers project to central and dorsolateral striatum. Coinciding with monkey anatomy, we also found a tripartite division of SN with similar anatomical and connectivity profiles (Figure 2B and Figure 2-Figure Supplement 1). Specifically, our mSNc corresponds to the monkey pars dorsalis and connects with ventral striatum; lSNc corresponds to the ventrolateral columnar part of SNc and connects to the motor regions of dorsal striatum. Finally, our vSN corresponds to the ventral densocellular portion that projects to the middle, associative, part of the striatum (Haber, 2014). Furthermore, the cortical projections of SN subdivisions we identified using diffusion tractography also fit with this limbic (mSNc), associative (vSN), and somatomotor (lSNc) organization (Figures 3-5). This specific association of lSNc with sensorimotor cortex explains its crucial role in the motor symptoms of Parkinson disease, where ventrolateral SNc is preferentially targeted by neurodegeneration (Gibb and Lees, 1991).

The inverted dorsal/ventral topography of SN to striatum projections (Haber, 2014) has also been described in human brain (Chowdhury et al., 2013). There, the authors used diffusion tractography in 30 individuals to parcellate SN based on anatomical connections with two targets in striatum. The dorsal SN mainly connected to ventral striatum, while the ventral SN preferentially projected to dorsal striatum. In contrast to this study, we used the whole-brain connectivity profiles to identify the subareas within SN instead of using predefined regions of interest restricted to striatal regions.

Recently, whole-brain tractography was also performed on the HCP dataset to identify the major brainstem white matter tracts (Meola et al., 2016). Two distinct fiber tracts were found projecting through SN: the frontopontine tract (FPT) connecting prefrontal cortex and anterior part of SN and running through the anterior limb of the internal capsule, and the corticospinal tract (CST) connecting motor cortex and posterior part of SN and passing through posterior limb of the internal capsule. This accords with our fiber tracking results, with vSN mainly connecting to the prefrontal cortex through the anterior limb of the internal capsule adjacent to the anterior dorsal striatum (including the body of caudate and anterior part of putamen), and lSNc preferentially connecting to the sensorimotor cortex via the posterior limb of the internal capsule and adjacent posterior dorsal striatum (including the tail of caudate and posterior putamen). We identified an additional fiber tract, i.e. a mesolimbic pathway connecting medial SNc with ventral striatum, vmPFC and OFC.

The somatomotor to associative to limbic (from lateral to medial) organization of SN accords with the cortical arrangement of information flow proposed by Mesulam (Mesulam, 1998), in which unimodal areas project to heteromodal associative, and then to prelimbic and limbic regions. A recent study proposed a similar gradient of cortical information processing based on resting state fMRI data from the HCP (Margulies et al., 2016). Our results suggest that the somatomotor to associative to limbic principle of cortical organization appears to be reflected in the SN.

### Value and Salience coding in SN projections

We found a double dissociation between coding of value and salience within SN subdivisions and their projections. Specifically, mSNc encoded monetary value in the gambling task, showing significantly greater BOLD response to wins than losses (Figure 6). Medial SNc preferentially connects to limbic areas including ventral striatum, ventral pallidum, hippocampus, amygdala and OFC/vmPFC (Figure 4). These brain regions have been reported to support value-based reinforcement learning (Garrison et al., 2013; Glimcher, 2011), and goal-directed behaviors (Goto and Grace, 2005), and have been implicated in drug addiction (Nutt et al., 2015). By contrast, vSN encoded salience, showing a similar BOLD response to rewarding and aversive events (Figure 6). VSN mainly connects to the prefrontal cortex and salience network, including lateral frontal cortex, dorsomedial prefrontal cortex, dACC and anterior insula (Figure 4). These brain areas are associated with attention, orientation and cognitive control (Menon and Uddin, 2010; Seeley et al., 2007; Uddin, 2015). Finally, the lSNc subdivision also appeared to encode salience, responding equally to monetary gains and losses. In contrast to the mesocortical pathway derived from vSN, the predominant projections of lSNc were with the motor cortex, premotor cortex, supplementary motor area, and posteriorly into the parietal cortex (Figure 4). The value/salience dissociation of lateral and medial SNc corresponds to the findings from recordings in Macaque midbrain dopamine neurons (Bromberg-Martin et al., 2010; Matsumoto and Hikosaka, 2009). A somewhat similar functional dissociation in SN was reported in a fMRI study with a Pavlovian learning paradigm (Pauli et al., 2015), with the lateral SN encoding a prediction signal for aversive events, and the medial SN encoding a reward prediction error signal for appetitive learning.

### SN and impulsivity

Impulsivity has been reported to contribute to a wide range of psychopathology including bipolar disorder (Swann et al., 2009), ADHD (Winstanley et al., 2006), alcohol and substance dependence (Ersche et al., 2010), pathological gambling (Leeman and Potenza, 2012), and addictive behaviors in Parkinson’s Disease (Averbeck et al., 2014; Dagher and Robbins, 2009). A current account of impulsivity assigns a key role to midbrain dopamine neurons, which modulate choice behaviors through the direct and indirect corticostriatal pathways (Buckholtz et al., 2010; Dalley and Roiser, 2012). Specifically, phasic bursts of dopamine firing enhance impulsive and risk-taking behaviors through D1 receptors within the direct pathway, while pauses in dopamine firing activate inhibitory control through D2 receptors within the indirect pathway (Collins and Frank, 2014; Cox et al., 2015). Our findings suggest that impulsivity may also be a reflection of top-down cortical and striatal control of SN activity.

The multi-dimensional view of impulsivity proposes at least two major components (Meda et al., 2009), namely impulsive action and impulsive choice. Here we included two different impulsivity measures: the Delay Discounting task (impulsive choice) and Flanker inhibitory control task (impulsive action), and explored the neural basis of these two constructs in terms of brain activity and connectivity.

Our results suggest that two different dopamine systems modulate these two components of impulsivity in parallel. Specifically, decisional impulsivity, measured by the Delay Discounting task, was associated with the value-coding system (Figure 7A) and mesolimbic pathways connecting mSNc, VS and vmPFC (Figure 8). Stronger value-coding signals in these areas or weaker inter-regional connectivity were associated with more impulsive choices during delay discounting, meaning higher preference for immediate and smaller rewards. The negative association between decisional impulsivity and mSNc-vmPFC connectivity might reflect inhibitory top-down control of mSNc dopamine signalling from vmPFC (Dalley et al., 2011). Additionally, reduced top-down control might be reflected in greater or maladaptive phasic dopamine response to rewards, as reflected here in greater BOLD response to wins versus losses. Evidence has been shown in primates that vmPFC can affect dopamine neuron activity indirectly via the nucleus accumbens (Haber and Knutson, 2010). Meanwhile, motor impulsivity measured by the Flanker inhibitory control task was associated with the salience-coding system (Figure 7B) and mesocortical pathways projecting between vSN and dACC and anterior insula (Figure 8). Stronger BOLD signals in the salience network or stronger cortico-striatal projections predicted better attentional inhibitory control. This finding coincides with the theory that anterior insula plays an important role in inhibitory control by increasing the saliency of stimuli, especially for unexpected events (Cai et al., 2014; Ghahremani et al., 2015).

## Limitations

We included a large population of healthy young subjects acquired from the public HCP dataset. Multimodal data included structural, diffusion-weighted, and functional MRI, as well as behavioural impulsivity measures. There were a few missing imaging or behavioral data and some datasets failing during additional preprocessing. The final dataset included 485 subjects for the gambling-task fMRI data, 430 subjects for the diffusion data, and 488 subjects for the behavioral measures. In the end, we had over 400 overlapping subjects who had all three modalities.

The SN is a small nucleus located in the brainstem, where MRI data usually suffer from distortions and signal losses. Partial volume effect might have impacted the imaging data, especially for fMRI. However, in the HCP data, these problems have been mitigated by advanced high-resolution imaging sequences and preprocessing (Glasser et al., 2013; Setsompop et al., 2013). Still, one potential limitation of the current study is inferring midbrain dopaminergic projections from diffusion tractography. Diffusion tractography has several known limitations, including the inability to perfectly resolve crossing fibers, a relatively high susceptibility to false positives and negatives and a tendency to terminate in gyral crowns as opposed to sulci, resulting in diminished anatomical accuracy (Jbabdi et al., 2015; Jones et al., 2013; Thomas et al., 2014). A greater concern however is the possibility of systematic biases in probabilistic tractography that may give an incorrect impression of whole-brain SN connectivity patterns. For example, it is accepted that connections are less likely to be detected if they travel a long distance, exhibit marked curvature or branching, travel close to cerebrospinal fluid, or pass through more complex white matter regions (Jbabdi et al., 2015). Several aspects of our results make this less likely. First, although the SN parcellation and projection maps were based on diffusion tractography, they also reveal a functional dissociation. That is, projection maps of SN subdivisions reflect a limbic, associative and somatomotor organization, rather than a purely geometric pattern. Moreover, our parcellation of SN accords closely with tract tracing studies in macaque (Haber, 2014). Another limitation is that connectivity measured by diffusion tractography cannot resolve the direction of a connection, which makes it impossible to distinguish efferent dopamine projections from top-down fronto-nigral or striato-nigral projections.

Thus, although our results are consistent with those of Matsumoto and Hikosaka in monkeys (Matsumoto and Hikosaka, 2009), in which there is a medial to lateral gradient for reward/ salience coding, we cannot attribute either connectivity or fMRI activation to dopamine neurons per se. The population of neurons in VTA and SN is heterogeneous and includes GABAergic and glutamatergic projection neurons and interneurons (Henny et al., 2012; Morales and Margolis, 2017). Despite a reward/salience dissociation found in SN activation and projections, our results do not contradict the theory that dopamine neurons in SN/VTA are predominantly excited by reward and reward prediction error (Cohen et al., 2012; Fiorillo, 2013). Indeed, a plausible explanation for our findings is that the correlations between diffusion tractography and impulsivity reflect top-down control of dopamine signaling. Thus, greater connectivity from vmPFC to mSNc would enable optimum coding of value, and reduced decisional impulsivity. Similarly, the relationship between stronger ACC connectivity to vSN and better control of motor impulsivity may reflect cortico-nigral top-down control over dopamine neuron activity to salient stimuli. It is also notable that the fMRI results demonstrated a salience response (i.e. to both wins and losses) throughout the striatum, consistent with recordings in monkeys, and that our connectivity findings support the theory that predominantly reward versus predominantly salience coding SN neurons belong to different brain networks (Bromberg-Martin et al., 2010).

## Conclusions

We subdivided the human SN into three subpopulations according to anatomical connectivity profiles, with a dorsal-ventral and lateral-medial arrangement. Our three-way partition of SN reveals multiple dopaminergic systems in human SN, showing a limbic, cognitive and motor arrangement, and encoding value and salience signals separately through distinct pathways. Corresponding to this connectional arrangement, we also found dissociable functional response during the gambling task and correlations with impulsivity measures. Specifically, mSNc was involved in the value-coding system and associated with impulsive choice, while vSN was involved in the salience-coding system and associated with response inhibition. Building on the traditional RPE-model of dopamine signaling (Schultz, 1998), our study provides evidence for the connectional and functional disassociations of midbrain dopamine neurons in humans, which encode motivational value and salience, possibly through different dopaminergic pathways. We also extended the current view on the role of dopamine in impulsivity by uncovering different neural substrates for decisional and motor impulsivity.

## Acknowledgments

This work was supported by funding from the Canadian Institutes for Health Research and the Natural Sciences and Engineering Research Council of Canada.

